# Cortical Spheroid Model for Studying the Effects of Ischemic Brain Injury

**DOI:** 10.1101/2021.10.16.464587

**Authors:** Rachel M. McLaughlin, Amanda Laguna, Ilayda Top, Christien Hernandez, Liane L. Livi, Liana Kramer, Samantha Zambuto, Diane Hoffman-Kim

## Abstract

Stroke is a devastating neurological disorder and a leading cause of death and long-term disability. Despite many decades of research, there are still very few therapeutic options for patients suffering from stroke or its consequences. This is partially due to the limitations of current research models, including traditional *in vitro* models which lack the three-dimensional (3D) architecture and cellular make-up of the *in vivo* brain. 3D spheroids derived from primary postnatal rat cortex provide an *in vivo*-relevant model containing a similar cellular composition to the native cortex and a cell-synthesized extracellular matrix. These spheroids are costeffective, highly reproducible, and can be produced in a high-throughput manner, making this model an ideal candidate for screening potential therapeutics. To study the cellular and molecular mechanisms of stroke in this model, spheroids were deprived of glucose, oxygen, or both oxygen and glucose for 24 hours. Both oxygen and oxygen-glucose deprived spheroids demonstrated many of the hallmarks of stroke, including a decrease in metabolism, an increase in neural dysfunction, and an increase in reactive astrocytes. Pretreatment of spheroids with the antioxidant agent N-acetylcysteine (NAC) mitigated the decrease in ATP seen after 24 hours of oxygen-glucose deprivation. Together, these results show the utility of our 3D cortical spheroid model for studying ischemic injury and its potential for screening stroke therapeutics.

**Significance Statement:** Those who survive after suffering a stroke often have long-term cognitive or physical disabilities. There is currently only one available therapeutic, tissue plasminogen activator (tPA), and it must be administered within a few hours after the onset of stroke. As stroke prevalence increases with our aging population, there is a growing need for therapies to mitigate or reverse the resulting brain damage. Three-dimensional (3D) culture systems have the potential to screen novel therapeutics more reliably than traditional *in vitro* models. Here we present a novel 3D cortical spheroid ischemia model which replicates many of the characteristics of stroke and has the potential to be an effective tool in therapeutic development.

## Introduction

Stroke is the leading cause of serious long-term disability and the fifth leading cause of death in the United States (Benjamin et al. 2019). Ischemic stroke, which is caused by the blockage of an artery in the brain, causes 87% of all strokes (Benjamin et al. 2019). Currently, tissue plasminogen activator (tPA) is the only FDA-approved therapeutic agent to treat patients with acute ischemic stroke and acts by assisting in clot breakdown (The National Institute of Neurological Disorders and Stroke rt-PA Stroke Study Group 1995). To be eligible for tPA therapy, patients must receive the treatment within 4.5 hours of stroke onset and must not have an increased risk of bleeding, as tPA increases the risk of brain hemorrhaging (Peña et al. 2017; Powers et al. 2019). Three-quarters of stroke patients arrive to the hospital after the treatment window, and less than 5% of patients wind up receiving tPA (American Heart Association 2017). Once a clot has been cleared, patients have few options to work toward the recovery of lost function beyond rehabilitative therapy and time. The overwhelming majority of patients are ineligible to receive thrombolytic therapy, highlighting the need for improved therapeutic options.

The scientific community has identified a number of cellular and molecular pathways that are characteristic of ischemic brain injury. Immediate mechanisms of damage include decreased intracellular pH, decreased ATP production, initiation of free radical production, and increased intracellular Na^+^ leading to membrane depolarization and excitotoxicity (Lipton 1999). Inflammation and programmed cell death follow, further contributing to brain tissue damage (Mergenthaler and Meisel 2012). While traditional stroke models have been informative, hundreds of therapies found successful in these models have failed to translate to humans (Barthels and Das 2020; Sommer 2017). Testing therapeutics in two-dimensional (2D) cell culture is cost-effective and high throughput but does not accurately represent *in vivo* tissue. Animal models are better at modeling tissue-drug interactions but are expensive, less reproducible, and time consuming to use. In comparison with 2D cell culture, three-dimensional (3D) spheroids are able to more accurately mimic the microenvironment and cell diversity found *in vivo*. It has been shown that cells cultured in 3D have more physiologically relevant cell morphologies and cell-cell interactions and are able to produce their own extracellular matrix (Jensen and Teng 2020).

We have previously reported a self-assembled, scaffold-free, three-dimensional *in vitro* cortical spheroid model. These spherical microtissues are derived from primary postnatal rodent cortex and have a cellular composition, tissue stiffness, and cell density similar to what is found in the *in vivo* cortex (Dingle et al. 2015). Neurons in these microtissues are electrically active and form synaptic connections within two weeks of seeding (Dingle et al. 2015). Hundreds of spheroids can be created from every rodent dissected, increasing the number of conditions that can be tested and reducing the number of animals needed for an experiment. The cortical spheroids contain a diverse set of brain cell types, including neurons, astrocytes, microglia, oligodendrocytes, and neural progenitor cells (Dingle et al. 2015). This model contains spontaneously formed capillary-like networks, which include many of the components that are essential to forming the neurovascular unit (Boutin et al. 2018). Previously we showed that these capillary-like networks are composed of endothelial cells surrounding a lumen, and these endothelial cells are associated with tight junctions. This is a unique feature of our model that is not yet replicated in other *in vitro* models, such as organoids. The size and cellular composition of the spheroids are very reproducible, making them an excellent candidate to use for high-throughput assays (Dingle et al. 2015).

In this study, cortical spheroids were deprived of oxygen and/or glucose for 24 hours to emulate the different components of ischemic injury. We evaluated several different molecular and cellular pathways that have been implicated in stroke-induced brain injury. We found a decrease in ATP production, presence of reactive astrocytes, loss of cellular structural integrity, and dysfunction of calcium dynamics after oxygen or oxygen-glucose deprivation. Our assessment of N-acetylcysteine (NAC) revealed that pretreatment with this antioxidant mitigated the loss of ATP after oxygen-glucose deprivation. Our results suggest that 3D cortical spheroids reproduce key features of ischemic injury and can be a useful model for therapeutic screening.

## Materials and Methods

### Cell isolation

All animal procedures were conducted in accordance with the guidelines established by the NIH and approved by Brown University’s Institutional Animal Care and Use Committee. Primary cortical tissue was isolated from postnatal day 0–2 CD rats (Charles River); anogenital distance was used to determine sex. Cells were isolated using a modified version of the protocol developed by BrainBits, as previously described (Dingle et al. 2015). Briefly, cortical dissection was performed in a buffer of Hibernate A (BrainBits) with 1X B27 supplement (Invitrogen), and 0.5 mM Gluta-Max (Invitrogen). Rat cortices were cut into small pieces and digested for 30 minutes at 30°C in 2 mg/mL papain dissolved in Hibernate A without Calcium (BrainBits). After removal of papain, tissues were triturated with a fire polished pipette in Hibernate A buffer solution. Cell suspension was filtered through a 40 μm cell strainer and subsequently centrifuged at 150 xG for 5 min. The pellet was then resuspended in complete cortical medium: NeurobasalA medium (Invitrogen) with 1X B27, 0.5 mM Gluta-Max, and 1X Penicillin-Streptomycin (Invitrogen). The suspension was then filtered, subsequently centrifuged, resuspended, and centrifuged again. Cell viability and count were determined using a trypan blue exclusion assay (Invitrogen). 2D viability control cultures were seeded in 12-well cell culture plates on wells coated with poly-D-lysine (Sigma-Aldrich) at a density of 288,000 cells/well.

### 3D spheroid culture

Autoclaved-sterilized powder UltraPure agarose (Invitrogen) was dissolved via heating to 2% wt/vol in phosphate-buffered saline (PBS) and poured into spheroid micromolds with 96 round pegs of 400 μm diameter (#24-96-Small, Microtissues, Inc) (Dingle et al. 2015). After cooling, agarose gels were removed from the molds and transferred to 24-well cell culture plates. Agarose gels were equilibrated in incomplete cortical medium (complete cortical media without B27) with 3 exchanges over the week prior to dissection. Gels were equilibrated in complete cortical media the day of dissection. For seeding, the appropriate number of cortical cells was resuspended in cortical medium at a volume of 75 μL per agarose gel. Equilibration medium was aspirated from gels and 75 μL of cell suspension was added to each agarose gel. Cortical cells were seeded at a density of 4000 or 8000 cells per microwell. Cortical cells were then allowed to settle into microwells for 60 min at 37°C with plates shaken every 20 minutes to guide cells in microwells, and once cells were settled, 1 mL of complete cortical medium was added per well. Cell culture medium was changed at 48 hours and every 2–3 days following.

### Ischemic injury

Spheroids were fed every 2-3 days as described previously. On the desired day *in vitro* (DIV), gels were placed under one of four conditions: control, glucose deprivation (GD), oxygen deprivation (OD), or oxygen-glucose deprivation (OGD), for a total of 24 hours. Control gels were washed twice with complete cortical media and placed directly in a 37°C incubator with room air and 5% CO_2_. GD gels were washed twice with glucose-free media and placed directly in a 37°C incubator with room air and 5% CO_2_. OD gels were washed twice with complete cortical media and placed in an anaerobic GasPak EZ container system (Beckton Dickison). OGD gels were washed twice with glucose-free media and placed in an anaerobic GasPak EZ container system (Beckton Dickison). Glucose free media consisted of Neurobasal-A without D-glucose and sodium pyruvate (Invitrogen), 1X B27 without antioxidants (Invitrogen), and 1X Penicillin-Streptomycin (Invitrogen). The GasPak EZ container system had an anaerobic sachet that sequestered oxygen and achieved O_2_ levels of less than 1% within 2.5 hours. Deprivation was started on DIV13 or 14 unless otherwise stated.

### ATP assay

Spheroids were gently pipetted from agarose gels. Individual spheroids in 100μL complete cortical media were transferred to an opaque-walled, clear bottom 96-well plate, using a 200μL pipette with the tip cut off to accommodate the spheroid. 100μL of CellTiter-Glo 3D Reagent (Promega Life Sciences) was added to each well. The plate was wrapped in aluminum foil and placed on a shaker for 10 minutes to induce lysis. The luminescence was measured using a Cytation5 (Biotek).

### Immunohistochemistry

After 24 hours of deprivation, spheroids were fixed by replacing media surrounding gels with a 4% paraformaldehyde (PFA)/8% sucrose solution overnight. The next day the gels were washed 3 times with PBS, leaving half an hour between each wash. The spheroids were then stored at 4°C until staining. All of the following steps were performed on a shaker at room temperature. The following antibodies were used: mouse anti-β-III-tubulin (Covance MMS-435P), rabbit anti-glial fibrillary acidic protein (GFAP, DAKO Z0334), rabbit anti-laminin (BTI BT594), mouse anti-MAP2 (Millipore MAB3418), Cy3 goat anti-mouse (Jackson 115-165-068), Alexa488 goat anti-rabbit (Jackson 115-545-146), and Cy3 goat anti-rabbit (Jackson 111-165-144). Spheroids were permeabilized and blocked with 3D Blocking Solution which consisted of 1% Triton X-100 (TX), 10% normal goat serum (NGS), and 4% bovine serum albumin (BSA) in PBS for 2 hours, and subsequently incubated in primary antibodies diluted in 3D Blocking Solution overnight. Spheroids underwent two 2-hour washes with 0.2% TX in PBS (PBT), followed by one 2-hour 3D Blocking Solution wash. Spheroids were incubated with secondary antibodies in 3D Blocking Solution overnight. Spheroids underwent two 2-hour PBT washes and incubated in 1 mg/mL 4’,6-diamidino-2-phenylindole (DAPI) in PBT for 1 hour and returned to PBS.

Laminin and GFAP-stained spheroids used for image analysis were cleared using an ultraFast Optical Clearing Method (FOCM) before imaging (Zhu et al. 2019). FOCM reagent was made by first dissolving 30% wt/vol urea and 20% wt/vol D-sorbitol in dimethyl sulfoxide on the shaker overnight. The following day, 5% wt/vol glycerol was added to the solution and was placed on the shaker for at least 10 minutes. The FOCM solution was then added to spheroids and samples were allowed to incubate for 10 minutes before being plated onto confocal dishes in FOCM. All other stained samples were imaged in PBS.

### Imaging

All samples were imaged using an Olympus FV3000RS (Olympus) laser scanning confocal microscope under the control of Olympus Fluoview FV3000 software. Fluorescent images were acquired using a 10X air objective (Olympus, UPlan Super Apochromat, NA 0.4, WD 3.1 mm) or 30X oil objective (Olympus, UPlan Super Apochromat, NA 1.05, WD 0.80 mm); and laser diode (405, 488, and 561 nm) laser lines. Imaging settings were consistent between samples for all conditions within the same experiment. Image stacks were converted to TIFF format and max z-projections were reconstructed in ImageJ.

### LIVE/DEAD assay

Cell viability was evaluated using a LIVE/DEAD Viability/Cytotoxicity Kit (Molecular Probes). Cortical spheroids were incubated at 37°C for 1 hour in complete cortical media containing 2μM calcein AM (green-live cells) and 4μM ethidium homodimer-1 (red - dead cells). After washing once with warm complete cortical media, the spheroids were then transferred to a confocal dish. Samples were kept at 37°C throughout the imaging process.

### Calcium imaging

Calcium imaging was performed as previously described (Sevetson, Theyel, and Hoffman-Kim 2021). Briefly, cortical spheroids were incubated for 25 minutes in complete cortical media containing 5.2μM Oregon Green BAPTA-1 AM (ThermoFisher) and 0.02% wt/vol Pluronic F-127 (ThermoFisher). After washing once with warm complete cortical media, the spheroids were then transferred to a confocal dish. Samples were kept at 37°C throughout the imaging process.

### Image Analysis

GFAP-stained spheroids were imaged using a 30X objective. Images were captured 1.5μm in depth apart and z-stacks captured the entire volume of each spheroid. GFAP-positive areas were analyzed in ImageJ using the 3D Object Counter plugin. The object size and intensity threshold filters were selected to include areas that contained astrocytes while excluding background pixels; these settings were kept consistent for all spheroids for all conditions. Average voxel intensity of the astrocytes was calculated by dividing the integrated density of the GFAP-positive area by the total volume of the GFAP-positive area for each spheroid.

Laminin-stained spheroids were imaged using a 10X objective to maximize the number of spheroids captured in one imaging session. Images were captured 5μm in depth apart and z-stacks captured the entire volume of each spheroid. Capillary-like networks (CLNs) were analyzed in ImageJ using the 3D Object Counter plugin. The object size and intensity threshold filters were selected to include groups of pixels that were part of the CLN while excluding background pixels and these settings were kept consistent for all spheroids for all conditions. Volume of the CLN for each spheroid was determined by the 3D Object Counter plugin and included all of the laminin-positive area as selected by the filter settings. To normalize for differences in spheroid size, volume of the CLN was divided by the total spheroid volume, which was quantified using the DAPI staining.

Analysis of calcium videos was performed using a custom MatLAB script developed in the lab (Sevetson, Theyel, and Hoffman-Kim 2021). Calcium videos were captured at 25 frames per second for 1 minute for each spheroid imaged. Image stacks were registered, and regions of interest (ROIs) were manually designated based on visible cell borders. Additional ROIs were circled to monitor whole-sphere activity and to check for background noise. Fluorescence over time was extracted and normalized to the first 500 frames for each ROI and saved with corresponding ROI topographical information. Binary active/inactive designations were made based on which cells displayed increased activity (at least a 10% increase in fluorescence intensity) between 0.05 Hz and 1 Hz. Fluorescence traces were de-noised with a 0.5 second gaussian filter prior to correlation calculations and individual trace plotting.

### N-acetylcysteine treatment

N-acetylcysteine (NAC) (Sigma-Aldrich) was used at 1mM or 5mM and was added to gels immediately before the induction of control or OGD conditions. Control + 1mM NAC and control + 5mM NAC were washed twice with complete cortical media containing 1mM or 5mM NAC respectively before being placed in a 37°C incubator with room air and 5% CO2. OGD + 1mM NAC and OGD + 5mM NAC were washed twice with glucose-free media containing 1mM or 5mM NAC respectively before being placed in an anaerobic GasPak EZ container system (Beckton Dickison). After 24 hours, samples were prepared for the ATP assay as previously described.

## Results

The aim of our study was to determine if the cortical spheroid model replicated the injury pathways that are seen in ischemic injury/stroke including depletion of ATP, cytoskeletal damage, neural dysfunction, reactive astrogliosis, and eventually cell death (Lipton 1999; Mehta 2014). We studied the effects of glucose deprivation (GD) and oxygen deprivation (OD) in addition to the combined oxygen-glucose deprivation (OGD) on the model to parse out how each component contributes to ischemic injury. We show that OD and OGD replicate many of the ischemic injury pathways. In addition, we used the newly developed platform to test the therapeutic potential of N-acetylcysteine (NAC). We found that pretreatment with 1 or 5mM of NAC was able to decrease the loss of ATP after oxygen-glucose deprivation. These results suggest that our model will be a useful tool for studying stroke and testing potential therapeutics.

### Viability decreased after oxygen and oxygen-glucose deprivation

Ischemic injury is characterized by a steep decrease in ATP levels followed by an increase in both necrosis and apoptosis (Lipton, 1999). The loss of ATP, along with other molecular changes that occur during ischemic injury, can lead to both acute and delayed cell death (Lipton, 1999). To detect changes in ATP production, we used the CellTiter-Glo 3D Assay (Promega Life Sciences), which lyses cells and produces a luminescent glow in the presence of ATP proportional to the ATP concentration in the solution. Spheroids were seeded at 4000 cells/microtissue and were exposed to control, GD, OD, or OGD conditions starting on DIV13 or 14. Relative ATP content was decreased by approximately half in OD (0.54±0.03) and OGD (0.46±0.04) conditions as compared to control (1.0±0.03) (Figure 1A, n=22 spheroids per condition total across three independent experiments, mean±SEM). There was a slight decrease in ATP content after GD (0.89±0.02) as compared to control, but this was less dramatic of an effect as seen after OD or OGD.

**Figure 1:**
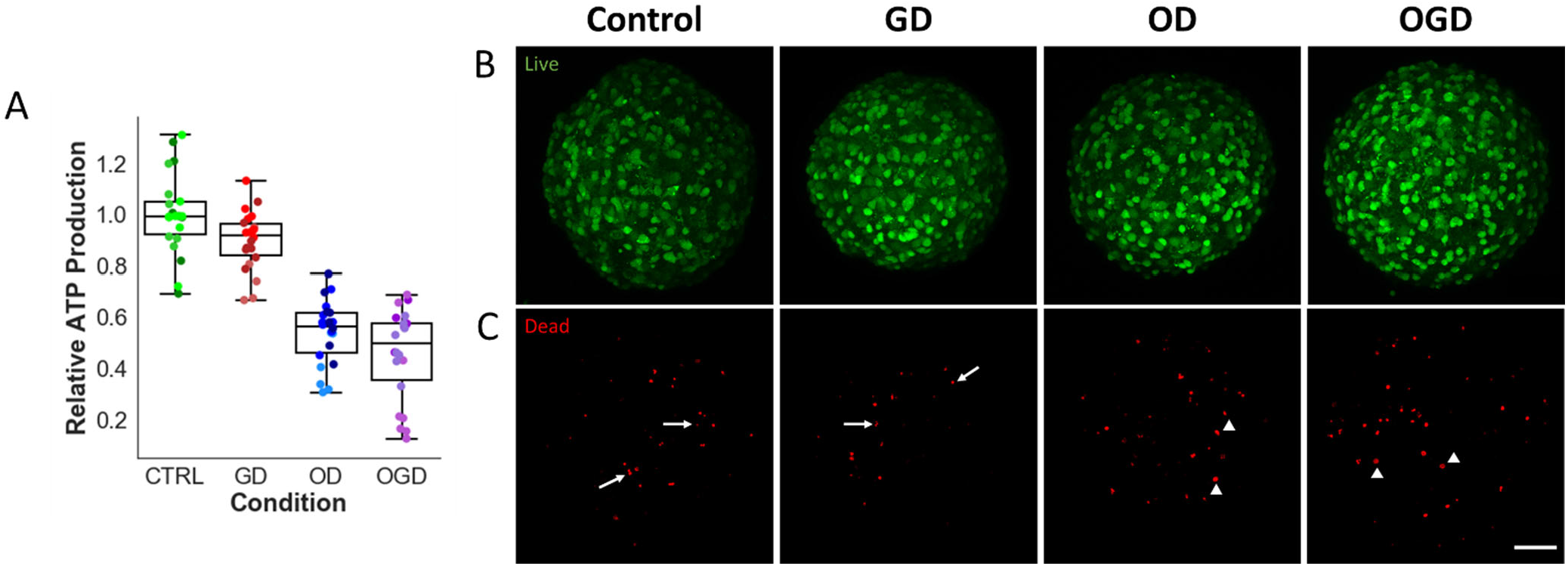
Cell viability decreased after oxygen and oxygen-glucose deprivation. Spheroids were exposed to glucose deprivation (GD), oxygen deprivation (OD), or oxygen-glucose deprivation (OGD) conditions for 24 hours. (A) Relative ATP production was decreased in OD and OGD conditions as compared to controls. Control (1.0±0.16), GD (0.89±0.11), OD (0.54±0.12) and OGD (0.46±0.18) mean ± SEM. n=22 spheroids per condition, 3 separate experiments represented by a different shade. Center lines of box plots represent median. Lower and upper whiskers represent minimum and maximum values of the datasets, respectively. Boxes extend from 25th to 75th percentiles. (B) Confocal z-projections of calcein AM or live cells and (C) ethidium homodimer 1 or dead cells. Arrows highlight smaller ethidium homodimer 1 positive cells in the control and GD conditions, arrowheads highlight larger ethidium homodimer 1 positive cells in the OD and OGD conditions. Scale bar is 50μm.

Ischemic injury causes cell death through several different mechanisms, and these mechanisms can be acute or delayed (Lipton 1999). It is thought that the delayed mechanisms of cell death may be reversible. To assess viability, we used calcein AM to label live cells green and ethidium homodimer 1 to label dead cells red. Although there was no observable difference in the number of dead cells between control and the three deprivation conditions, the ethidium homodimer 1 labeled cells appeared rounder and larger than those in control and GD conditions (Figure 1C). This suggests that although there was a decrease in cell metabolism, cell death in this model may be delayed.

### Neural cytoskeletal structure was disrupted following oxygen and oxygen-glucose deprivation

Cytoskeletal molecules play an essential role in maintaining cellular structure and mediating transport to and from dendrites and axons. Ischemic injury causes microtubule disruption and disorganization, which is often irreversible and leads to cell death (Lipton 1999). Reduction in microtubule associated protein 2 (MAP2) immunostaining is a reliable marker for irreversible neural damage after ischemic injury (Kharlamov et al. 2009). Axonal cytoskeletal breakdown after ischemia is characterized by beaded neurofilament staining which represents failure of regular axonal neurofilament distribution (Hinman 2014).

To characterize changes in neural cytoskeletal structure after GD, OD, and OGD, we stained spheroids with MAP2 and beta-iii-tubulin. MAP2 is associated with microtubule assembly and is enriched in the dendrites of neurons. Beta-iii-tubulin is a structural component of microtubule networks specifically localized to neurons. Spheroids seeded at 4000 cells/microtissue were subjected to 24 hours of deprivation starting on DIV14 and were fixed on DIV15. Confocal z-projections of MAP2 and beta-iii-tubulin showed distinct cytoskeletal changes in OD and OGD treatments as compared to controls. MAP2 staining showed increased puncta and fragmented strands in OD and OGD conditions as compared to the long, continuous dendrites seen in control and GD (Figure 2A and B). Beta-iii-tubulin showed higher prevalence of puncta in OD and OGD conditions (Figure 2C and D). These results together indicate that cytoskeletal structure of neurons are disrupted after ischemic injury, mimicking what other stroke models have shown (Lipton 1999).

**Figure 2:**
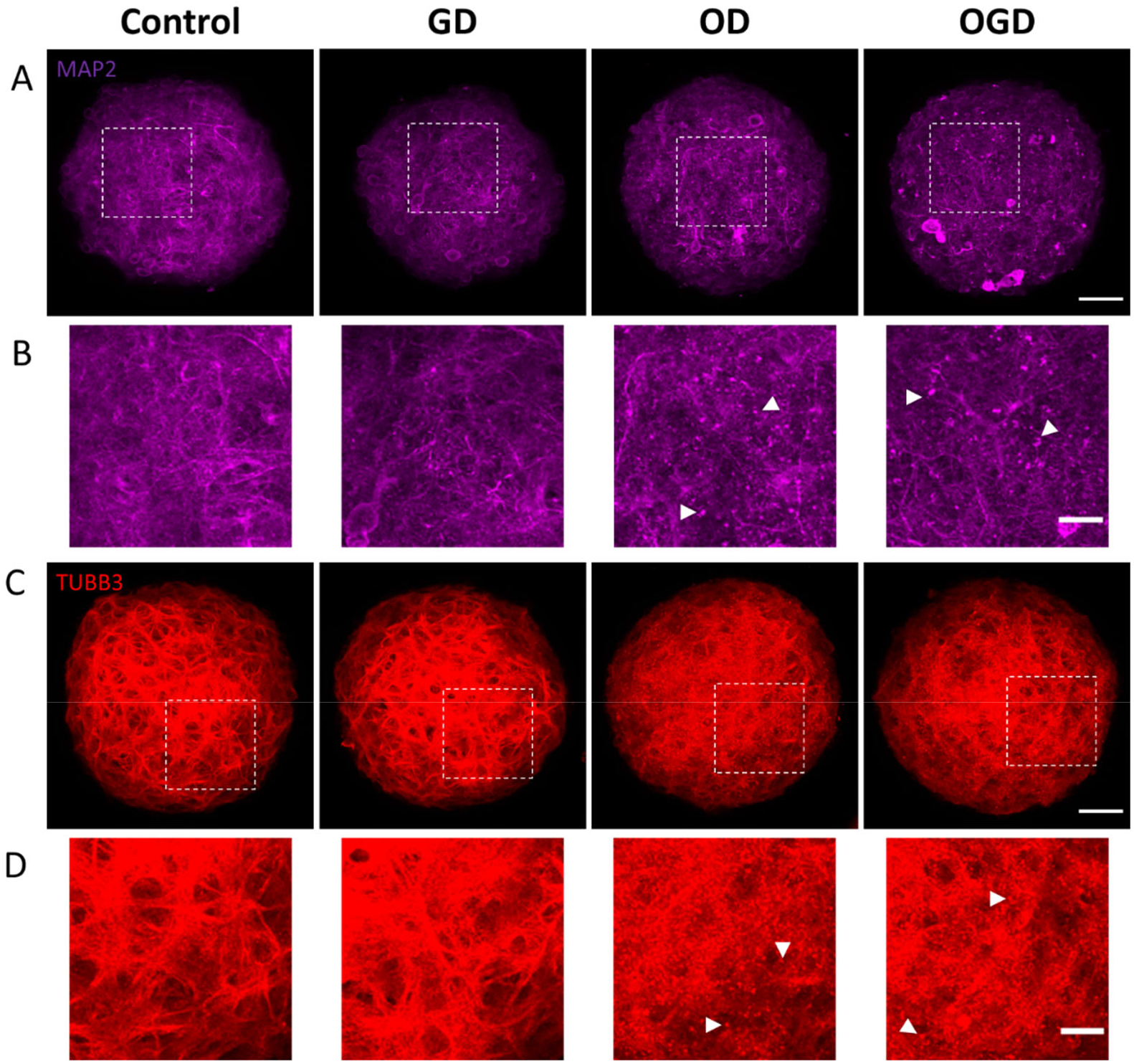
Neural cytoskeletal structure was disrupted following oxygen and oxygen-glucose deprivation. Spheroids were exposed to glucose deprivation (GD), oxygen deprivation (OD), or oxygen-glucose deprivation (OGD) conditions for 24 hours. (A) Confocal z-projections of whole spheroids stained for MAP2 and (B) enlargements of the squares in images directly above. Arrowheads highlight puncta in OD and OGD conditions. (C) Confocal z-projections of whole spheroids stained for beta-iii-tubulin and (D) enlargements of the squares in images directly above. Arrowheads highlight puncta in OD and OGD conditions. Scale bars are 50μm (A,C) or 20μm (B,D).

### Spheroids contained reactive astrocytes following oxygen and oxygen-glucose deprivation

Glial cells, such as astrocytes, are critical components of the central nervous system and regulate the neuroinflammatory response after stroke. Glutamate excitotoxicity during stroke induces reactive astrogliosis which is characterized by astrocyte proliferation, hypertrophy, and increased expression of glial fibrillary protein (GFAP) (Becerra-Calixto and Cardona-Gómez 2017; Sims and Yew 2017; Liu and Chopp 2016; Xu et al. 2020). Astrocytes can limit brain damage, reduce neuroinflammation, and are critical for blood-brain-barrier reconstruction after ischemic injury, but chronic reactive astrogliosis can hinder functional recovery and axon regeneration (Xu, 2020). Better understanding the role astrocytes play after ischemic injury could lead to the development of therapeutics targeting this inflammatory pathway. We therefore investigated whether the astrocytes in cortical microtissues showed signs of astrogliosis after GD, OD, or OGD. Spheroids seeded at 4000 cells/microtissue were subjected to 24 hours of deprivation starting on DIV14 and were fixed on DIV15. Confocal z-projections showed enhanced GFAP expression and increased cell soma width after 24 hours of OD and OGD as compared to control, with no substantial changes in astrocyte morphology after 24 hours of GD (Figure 3A and B). The average voxel intensity of the GFAP-positive astrocytes in the spheroids was higher in the OD (420.06±7.08) and OGD (444.95±6.52) conditions as compared to control (316.85±6.00) and GD (338.83±3.48) conditions (Figure 3C, n=8 spheroids per condition, mean±SEM).

**Figure 3:**
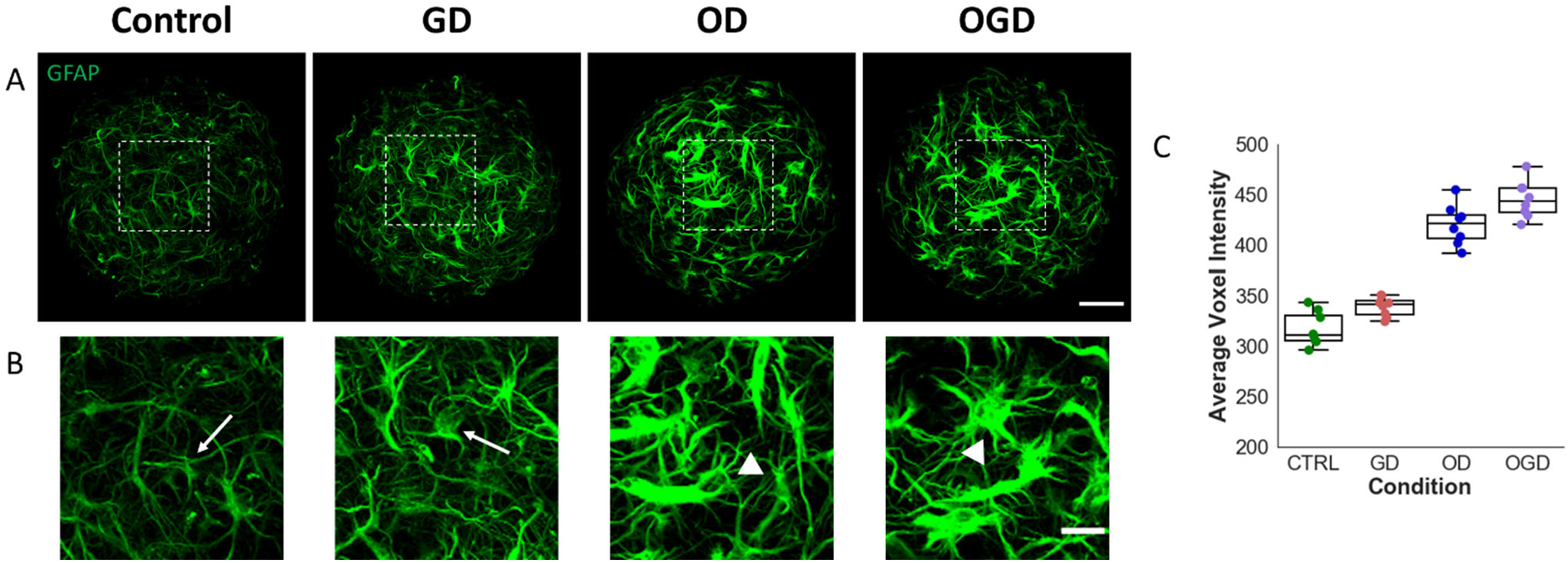
Spheroids contained reactive astrocytes following oxygen and oxygen-glucose deprivation. Spheroids were exposed to glucose deprivation (GD), oxygen deprivation (OD), or oxygen-glucose deprivation (OGD) conditions for 24 hours. (A) Top row images are confocal z-projections of whole spheroids stained for GFAP and (B) bottom row images are enlargements of the squares in the top row images. Arrows highlight phenotypically normal astrocytes in the control and GD conditions and arrowheads highlight hypertrophic astrocytes in the OD and OGD conditions. Scale bars 50 μm for the top row images (A) and 20 μm for the bottom row images (B). (C) Average voxel intensity was increased in OD and OGD conditions as compared to controls. Control (316.85±6.00), GD (338.83±3.48), OD (420.06±7.08), and OGD (444.95±6.52) mean ± SEM. n=8 per condition. Center lines of box plots represent median. Lower and upper whiskers represent minimum and maximum values of the datasets, respectively. Boxes extend from 25th to 75th percentiles.

### Spheroids showed capillary-like breakdown following oxygen and oxygen-glucose deprivation

Neurovascular function is linked to many CNS pathologies, including stroke. Breakdown of the bloodbrain barrier occurs early in pathogenesis and has been linked with more severe cognitive outcome (Hu et al. 2017). Ischemic injury and stroke induces endothelial cell dysfunction, tight junction disruption, and breakdown of the basement membrane, increasing the permeability of the vessel (M. Q. Jiang et al. 2017; Freitas-Andrade, Raman-Nair, and Lacoste 2020). As described earlier, cortical spheroids spontaneously form capillary-like networks (CLNs) that can be visualized using laminin staining (Boutin et al. 2018). We investigated whether there were structural changes in the CLNs after 24 hours of GD, OD, and OGD. Spheroids seeded at 8000 cells/microtissue were subjected to 24 hours of deprivation starting on DIV3 and were fixed on DIV4. This seeding density and timepoint were selected to optimize for the presence of CLNs, which are most abundant in the cortical spheroid model within the first week after seeding (Boutin et al. 2018). Confocal z-projections showed a disruption of CLN structure after 24 hours of OD and OGD as compared to control. CLNs in the OD and OGD conditions were more condensed and had less elongated and complex structures as compared to control and GD conditions (Figure 4A). These structural changes were quantified by measuring the volume of the laminin-positive structures and normalizing the CLN volumes to the volumes of the spheroids. CLNs in OD (0.31±0.04%) and OGD (0.33±0.04%) conditions were a much smaller percentage of total spheroid volume compared to the CLNs in control (3.74±0.36%) and GD (2.52±0.18%) conditions (Figure 4B, n=15 spheroids per condition, mean±SEM).

**Figure 4:**
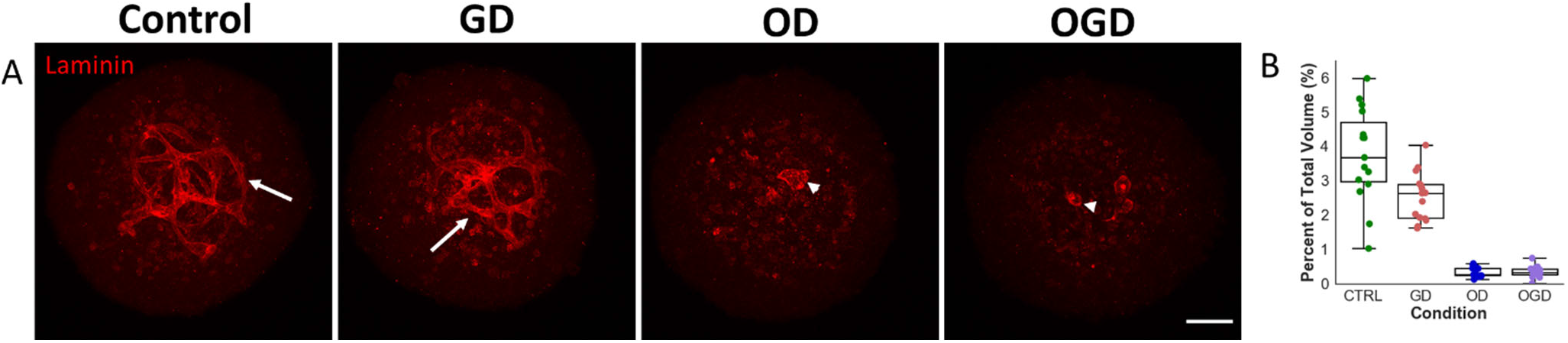
Spheroids showed capillary-like breakdown following oxygen and oxygen-glucose deprivation. Spheroids seeded at 8000 cells per microtissue were exposed to glucose deprivation (GD), oxygen deprivation (OD), or oxygen-glucose deprivation (OGD) for 24 hours starting on DIV3. Spheroids were fixed on DIV4. (A) Confocal z-projections of laminin structures show the extent of capillary-like networks (CLNs) in our spheroids. Arrows show complex tubular structures in the control and GD conditions, and arrowheads highlight condensed structures in the OD and OGD conditions. (B) The volume of the CLNs were normalized to the total volume of the spheroids and are presented as percent of total volume taken up by the CLN in each spheroid. The normalized CLN volume was decreased in OD and OGD conditions as compared to controls. Control (3.74±0.35%), GD (2.52±0.18%), OD (0.31±0.04%), and OGD (0.33±0.04%) mean ± SEM. n=15 per condition. Center lines of box plots represent median. Lower and upper whiskers represent minimum and maximum values of the datasets, respectively. Box extends from 25th to 75th percentiles.

### Calcium dynamics were disrupted after oxygen and oxygen-glucose deprivation

Within minutes of the onset of ischemic injury, there is a massive increase in intracellular Na^+^, widespread depolarization, and excitotoxicity, leading to synaptic dysfunction and neuronal cell death (Lipton 1999). When neurons are active, they influx calcium before releasing neurotransmitters, making detecting an increase in intracellular calcium a good proxy for detecting neural activity. We have previously described calcium dynamics in our model, which shows synchronized activity at DIV9 (Sevetson, Theyel, and Hoffman-Kim 2021). To see if this synchronized activity was disrupted by ischemic injury, we subjected spheroids that were initially seeded at 8000 cells/microtissue to GD, OD, or OGD at DIV8 and imaged the spheroids on DIV9 after 24 hours of deprivation. Sample calcium traces showed that control and GD spheroids had more cells with regular and synchronized calcium activity as compared to OD and OGD spheroids (Figure 5A). Quantification of these dynamic changes showed a decrease in the number of active cells in OD (16.8±3.1%) and OGD (10.6±2.6%) conditions as compared to control (61.0±5.3%) and GD (45.3±7.6%) conditions (Figure 5B, n=12 spheroids across two independent experiments, mean±SEM). There was very little correlation of active cells in OD (0.02±0.01) and OGD (0.01±0.01) conditions as compared to control (0.44±0.03) and GD (0.31±0.04) conditions (Figure 5C, n=12 spheroids across two independent experiments, mean±SEM).

**Figure 5:**
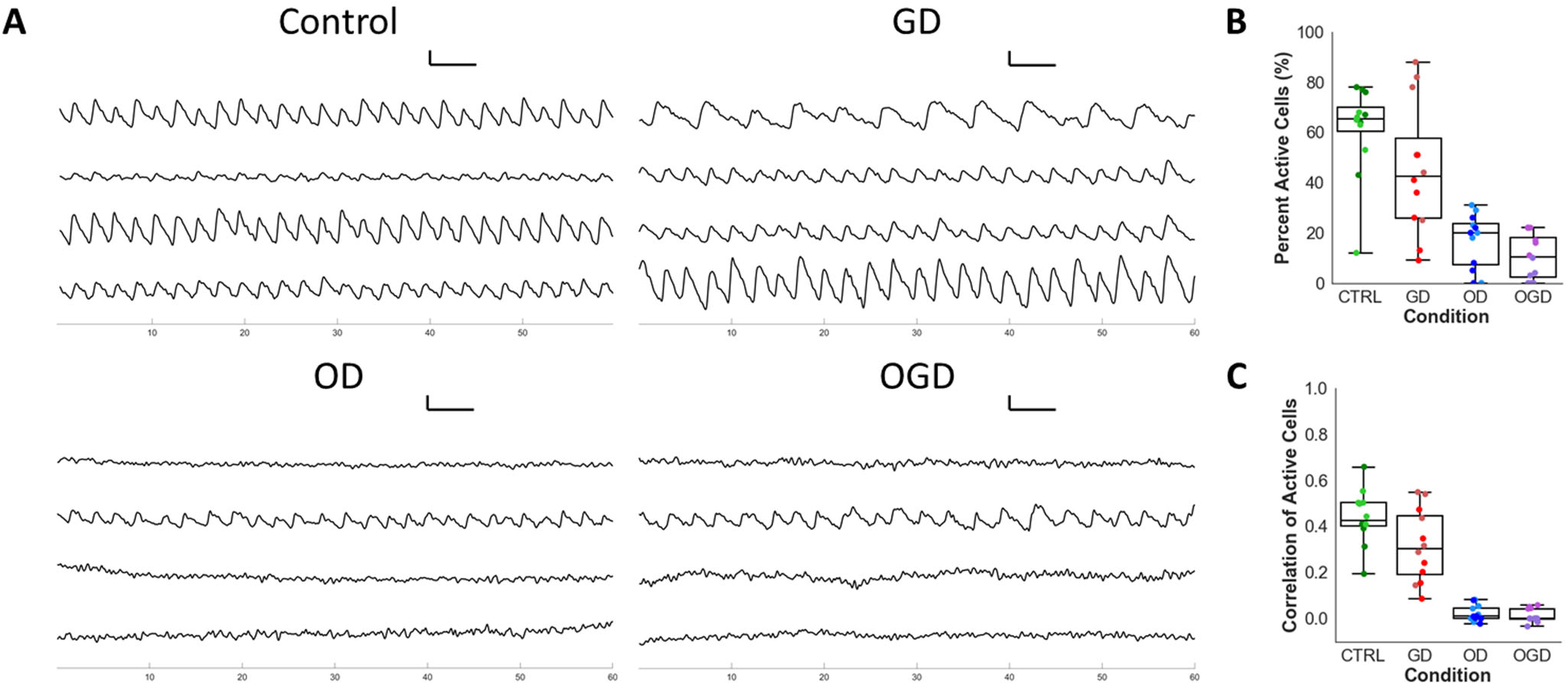
Calcium dynamics were disrupted after oxygen and oxygen-glucose deprivation. Spheroids seeded at 8000 cells per microtissue underwent 24 hours of glucose deprivation (GD), oxygen deprivation (OD), or oxygen-glucose deprivation (OGD) starting on DIV 8. On DIV 9, spheroids were incubated with Oregon Green Bapta-1 AM and imaged. (A) Sample line traces showing change in fluorescence over time. Control and GD spheroids have regular oscillatory patterns, while this activity is disrupted in the OD and OGD conditions. (B) Percent of active cells was decreased in OD and OGD conditions as compared to controls. Control (61.0±5.3%), GD (45.3±7.6%), OD (16.8±3.1%), and OGD (10.6±2.6%) mean ± SEM. (C) Correlation of active cells was decreased in OD and OGD conditions as compared to controls. Control (0.44±0.03), GD (0.31±0.04), OD (0.02±0.01), and OGD (0.01±0.01) mean ± SEM. n = 12 spheroids per condition, 2 separate experiments represented by a different shade. Center lines of box plots represent median. Lower and upper whiskers represent minimum and maximum values of the datasets respectively. Box extends from 25th to 75th percentiles.

### Pretreatment with N-acetylcysteine reduced loss of ATP content after oxygen-glucose deprivation

N-acetylcysteine (NAC) is an antioxidant that has been found to partially reduce the damage caused by ischemic injury and stroke in animal models (Z. Zhang et al. 2014; Turkmen et al. 2016). To see if NAC would mitigate the damage caused by OGD in our model, we treated spheroids seeded at 4000 cell/microtissue with 1mM or 5mM NAC at DIV13, immediately before the induction of control or OGD conditions. After 24 hours of deprivation, ATP production was measured. OGD spheroids pretreated with 1mM (0.60±0.05) or 5mM (0.72±0.04) had a higher relative ATP content than OGD spheroids that did not receive NAC treatment (0.45±0.02), with OGD spheroids pretreated with 5mM NAC obtaining ATP levels about halfway between OGD and control (1.0±0.04) conditions (Figure 6, n=8 spheroids per condition, mean±SEM).

**Figure 6:**
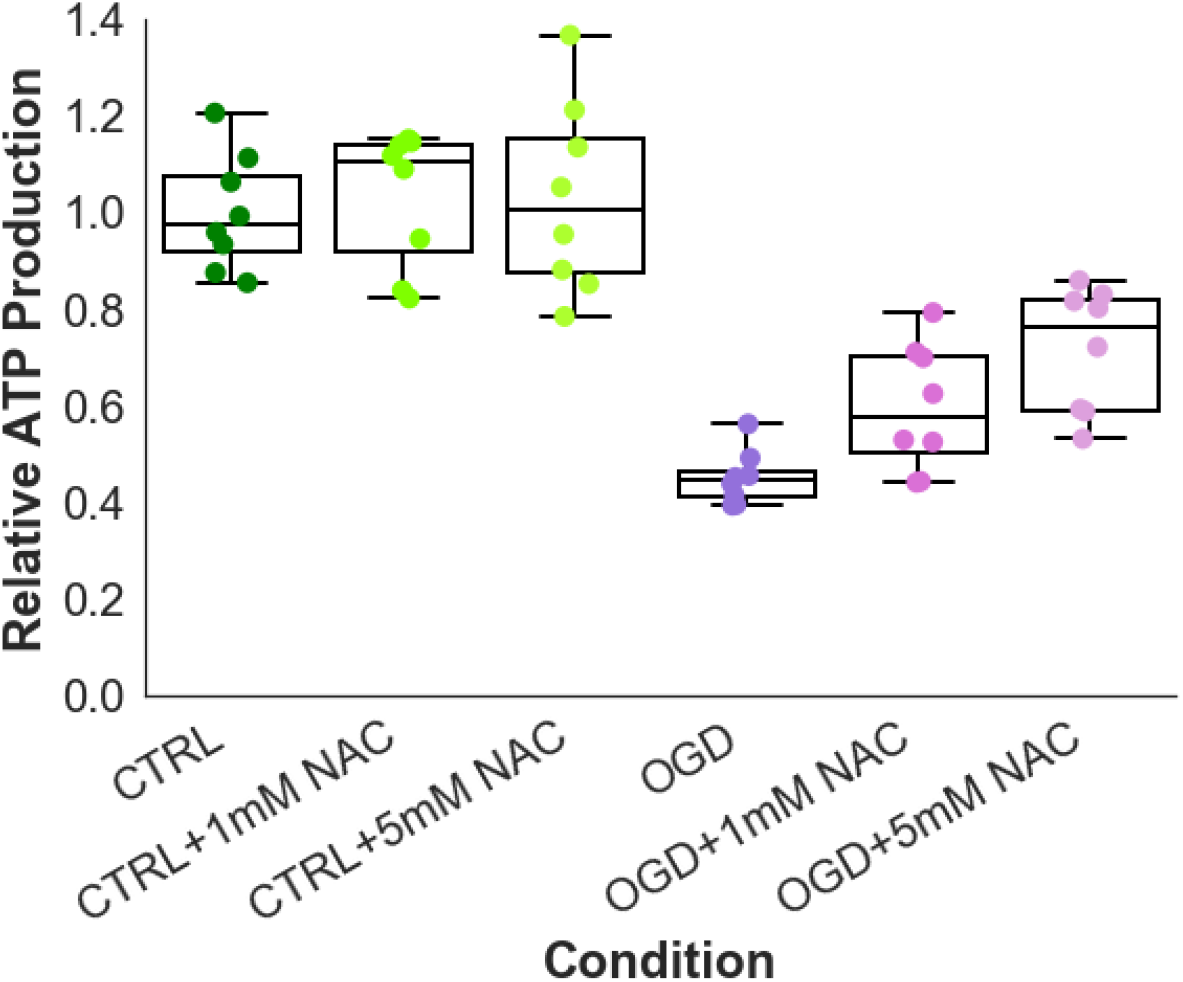
Pretreatment with N-acetylcysteine reduced loss of ATP production after oxygen-glucose deprivation. Spheroids were exposed to control, control plus 1mM of NAC, control plus 5mM of NAC, oxygen-glucose deprivation (OGD), OGD plus 1mM of NAC, or OGD plus 5mM of NAC for 24 hours. Pretreatment with NAC increased relative ATP production in OGD spheroids. Control (1.0±0.04), control plus 1mM NAC (1.03±0.05), control plus 5mM NAC (1.03±0.07), OGD (0.45±0.02), OGD plus 1mM NAC (0.60±0.05), control plus 5mM NAC (0.72±0.04), mean ± SEM. n=8 spheroids per condition. Center lines of box plots represent median. Lower and upper whiskers represent minimum and maximum values, respectively. Boxes extend from 25th to 75th percentiles.

## Discussion

Stroke is a devastating medical condition with few therapeutic options. Globally, about one in every four adults will suffer from a stroke in their lifetime, and about half of all stroke survivors have long-term disabilities (Virani et al. 2020; Donkor 2018). Despite decades of research and many different *in vivo* and *in vitro* models of ischemic stroke, hundreds of potential therapies have failed to translate to patients (Barthels and Das 2020; Sommer 2017). In this study, we presented a novel *in vitro* 3D cortical microtissue model for studying ischemic brain injury and testing potential stroke therapeutics. 3D self-assembled spheroids composed of primary postnatal rat cortical cells have the cellular composition, tissue stiffness, and cell density similar to the *in vivo* brain (Dingle et al. 2015). These spheroids contain complex cellular units important to brain function, including neural networks and capillary-like networks (Boutin et al. 2018; Sevetson, Theyel, and Hoffman-Kim 2021). Hundreds of spheroids can be produced per animal and spheroid production does not require expensive or proprietary reagents or equipment, making replication of our experiments relatively easy and cost-effective. This ischemic injury model provides an *in vivo*-relevant, high-throughput compatible, flexible platform to study the mechanisms of ischemic injury.

In this study, we used primarily female derived cortical spheroids to study the impact of ischemic injury. Females have a higher lifetime risk of stroke (1 in 5 females compared to 1 in 6 males) and often have greater disability from stroke than males (Virani et al. 2020). Despite these sex differences, males are massively overrepresented in both *in vitro* and *in vivo* preclinical trials, and many researchers do not know or do not report the sex of their cell sources (Karp and Reavey 2019). The use of male animals is about five-fold higher in both cardiovascular and neuroscience studies, and although some steps have been taken to improve the representation of female animals in preclinical research there is still huge room for improvement (Ramirez et al. 2017; Beery and Zucker 2011; Plevkova et al. 2021). There is a need to study ischemic brain injury and test potential stroke therapeutics in both female and male derived *in vitro* models, as both stroke outcome and drug interaction are sexually dimorphic (Virani et al. 2020; Karp and Reavey 2019). While this study focused on female derived microtissues, observations of male derived spheroids show similar phenotypic changes as those seen in female spheroids after OD and OGD (Supplemental Figure 1).

To model ischemic stroke, we induced OGD, a widely used paradigm for modeling ischemia *in vitro*. To achieve anaerobic conditions, we utilized a GasPak EZ container system that contained an anaerobic sachet. Similar systems have been used to investigate the effects of oxygen-glucose deprivation on *in vitro* blood-brain barrier permeability and have been used in a variety of other hypoxia models (Hind, England, and O’Sullivan 2016; Potapova et al. 2008; Sandhu et al. 2003; W.-H. Zhang et al. 2003). We additionally looked at GD and OD independently to determine how they contribute to ischemic injury. Brain ischemia leads to acidosis, oxidative stress, excitotoxicity, and eventually inflammation and cell death (Lipton 1999). In our study, we identified several different molecular, cellular, and functional changes that occur in this model after OD or OGD. Interestingly, GD alone had less of a detrimental impact on cortical spheroid health. Though it is possible that this may be due to the hyperglycemic nature of the cell culture environment, our results indicate that lack of oxygen is more detrimental to brain tissue than lack of glucose. After 24 hours of OD or OGD, there was a significant decrease in ATP production. A decrease in ATP occurs when there are fewer metabolically active cells, which may be due to lower metabolism in a live cell population or a decrease in the number of living cells. To determine whether the decrease in ATP corresponded to an increase in cell death, we performed a LIVE/DEAD assay. Our results indicated that there was no significant increase in the number of ethidium homodimer 1 positive, or dead, cells after 24 hours of deprivation. In other models, mitochondria depolarization and decrease in cellular metabolism after ischemia precedes cell death (Lipton 1999). This suggests that this 24-hour OGD paradigm models a the critical timepoint where there is significant cell damage, but there are still viable cells that can potentially be rescued.

Astrocytes are the most abundant glial cell in the brain and have a diverse set of important functions including synapse modulation and maintaining ion homeostasis (Liu and Chopp 2016). After ischemic brain injury astrocytes become reactive and upregulate glial fibrillary acidic protein (GFAP) (Becerra-Calixto and Cardona-Gómez 2017). The morphology of the astrocyte changes with the upregulation of GFAP, the cell body enlarges and the processes become thicker and bushier (Sims and Yew 2017). In the cortical spheroid model, astrocytes in the OD and OGD conditions have enlarged cell bodies, thicker processes, and have a higher expression of GFAP, indicating they have undergone astrogliosis. This replicates what has been observed in other stroke models. Reactive astrocytes can either exacerbate cytotoxicity or can help prevent further damage after injury, though the regulation of these dual roles is not completely understood (Xu et al. 2020). Astrocytes are increasingly becoming recognized as potential therapeutic targets after stroke or other brain injuries and modulating their ability to mitigate damage after injury could have huge implications on the recovery of stroke patients.

Neurovascular unit dysfunction is another major outcome of stroke. Ischemic injury induces endothelial cell dysfunction and tight junction disruption, increasing the permeability of blood vessels (X. Jiang et al. 2018; Freitas-Andrade, Raman-Nair, and Lacoste 2020). Proteases secreted from microglia break down the basement membrane surrounding the vessels, which leads to more vessel leakage (Freitas-Andrade, Raman-Nair, and Lacoste 2020). For the first week *in vitro*, cortical spheroids contain spontaneously formed capillary-like networks (CLNs). These networks contain endothelial cells with lumens that are surrounded by basement membrane proteins as well as other key cells of the NVU (Boutin et al. 2018). Most *in vitro* models of the NVU lack essential cell types or have an inside-out orientation with endothelial cells ensheathing brain tissue (Bhalerao et al. 2020). CLNs spontaneously form inside of the cortical spheroids, mimicking the *in vivo* structure more faithfully. These CLNs are associated with basement membrane proteins and can be visualized using a laminin stain (Boutin et al. 2018). In our study, we found that after OD or OGD there was a significant disruption in laminin structure. After OD or OGD, the total volume was significantly decreased, and the laminin-positive structure became less elongated and complex, and more condensed and spherical. This change in the laminin staining suggests that complex CLN structure collapsed after OD and OGD in this model. Future interventions that are found to prevent this structural collapse after ischemic injury may be useful for repairing or preventing neurovascular damage *in vivo*.

Neuronal injury after ischemia is complex and multifaceted (Lipton 1999). Cytoskeletal structure damage and loss of synaptic transmission occurs early after ischemic injury (Lipton 1999). In our study, we show that there is substantial cytoskeletal damage in both the dendrites and the axons after OD or OGD. Both MAP2 and beta-III-tubulin staining show the loss of strand-like structures and an increase in punctate or aggregated proteins. Axon and dendritic beading and fragmentation are signs of axon degeneration and occur after excitotoxicity (King et al. 2013). Neuronal cytoskeletal loss is thought to be a sign of the transition from reversible to irreversible cellular damage, making it an exciting potential therapeutic target (Mages et al. 2018). In addition to cytoskeletal changes, we observed a significant disruption in neural network dynamics after OD or OGD. We used calcium imaging to observe neural network changes after deprivation. At nine days *in vitro*, there are synchronous oscillatory calcium events in our model (Sevetson, Theyel, and Hoffman-Kim 2021). After OD or OGD, there are fewer active cells and the calcium activity of the active cells is less synchronized. This indicates that OD and OGD disrupt neural activity and communication between cells. Future studies will explore the mechanisms of this disruption.

NAC is a compound with established antioxidant and anti-inflammatory properties. It has been shown in several animal studies to be useful in reducing the damage caused by ischemic injury (Z. Zhang et al. 2014; Turkmen et al. 2016). Of note, as a promising treatment for stroke, it is most effective as a pretreatment or as a very early intervention (Tardiolo, Bramanti, and Mazzon 2018). Pretreatment of cortical spheroids with NAC was able to partially rescue the loss of ATP production seen after OGD (Figure 6). This result indicate that our model is able to show the beneficial effects of a potential stroke intervention.

Our cortical microtissue model suitably mimics neuropathologies such as ischemic injury and will allow for *in vitro* testing and development of novel therapeutics. Interestingly, our model only shows a slight increase in dead cells after 24 hours of deprivation, indicating that therapeutic interventions could be tested in our model to prevent or reverse damage before widespread permanent cell death. These data together show that our 3D *in vitro* cortical spheroid model replicates many of the hallmarks of ischemic injury and imply the potential for cortical spheroids to become a powerful screening tool for potential stroke therapeutics.

## Author Contributions

RMM and DHK designed the research. RMM, IT, and LLL conducted research. LK and SZ performed unpublished preliminary experiments. RMM, AL, and CH analyzed data. RMM and DHK wrote the paper.

## Acknowledgements

The authors thank Harrison Katz and Revanna Navarro for assistance with data analysis and Jess Sevetson for technical assistance and helpful discussion.

---

**Supplemental Figure 1:**
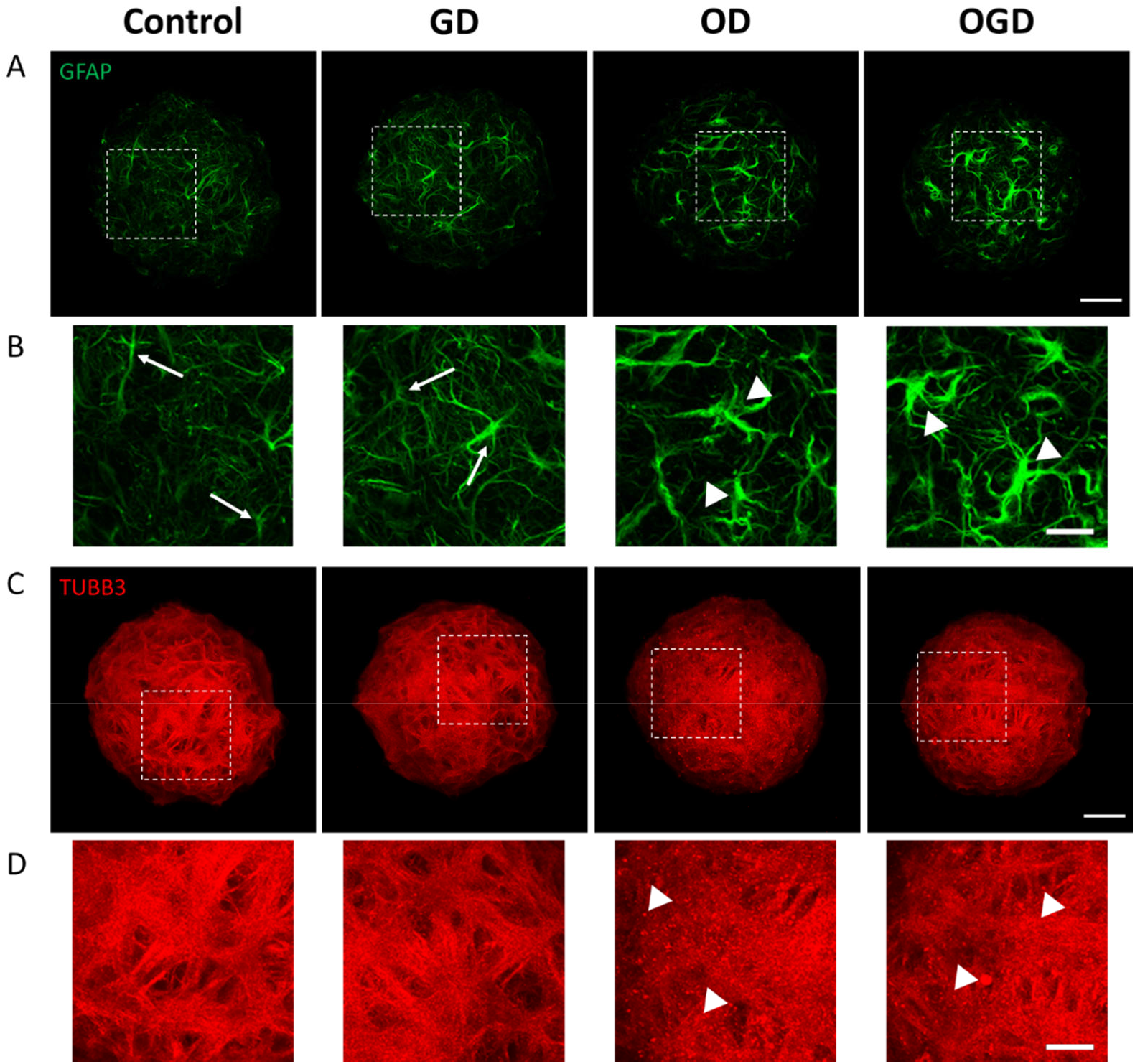
Male derived spheroids showed similar phenotypic changes after oxygen and/or glucose deprivation as female derived spheroids. Spheroids were exposed to glucose deprivation (GD), oxygen deprivation (OD), or oxygen-glucose deprivation (OGD) conditions starting on DIV14 for 24 hours. (A) Confocal z-projections of whole spheroids stained for GFAP and (B) enlargements of the squares in images directly above. Arrows highlight phenotypically normal astrocytes in the control and GD conditions and arrowheads highlight hypertrophic astrocytes in the OD and OGD conditions. Scale bars 50 μm (A) or 20 μm (B). (C) Confocal z-projections of whole spheroids stained for beta-iii-tubulin and (D) enlargements of the squares in images directly above. Arrowheads highlight puncta in OD and OGD conditions. Scale bars are 50μm (C) or 20μm (D).

